# Deficits in forelimb reach learning in a mouse model of Fragile X syndrome

**DOI:** 10.1101/2024.11.20.623831

**Authors:** Leanne F. Young, Ann Derham, Rui Zhu, Aparna Suvrathan

**Author notes:** These authors contributed equally (Ann Derham, Rui Zhu).

## Abstract

Fragile X syndrome is a leading cause of intellectual disability and autism spectrum disorder, for which therapies are limited. A mouse model of Fragile X syndrome, the *Fmr1* knockout (KO) mouse, has been particularly valuable for interrogating the molecular, cellular, and circuit mechanisms that underlie the neurological deficits seen in this syndrome. Key deficits in Fragile X syndrome include impairments in social behaviors, cognition, and motor learning. Given the difficulties in extrapolating more complex human behaviors to mouse models, simple motor behaviors are a particularly tractable form of learning to study in the mouse. We investigated a form of forelimb reach learning in *Fmr1* KO mice, precisely quantifying different parameters of the task using both manual analysis and DeepLabCut-based tracking of reach trajectories. While *Fmr1* KO mice show impaired learning overall, our results demonstrated that the presence or absence of a cue that signals reward alleviates some of the deficits. In addition to a single metric of success in learning, we determined the specific parameters of the motor behavior that were responsible for that success or failure. In particular, our results suggested that *Fmr1* KO mice showed impaired improvement in the trajectory of the reach, reflected by a greater likelihood of completely missing the target, and in a lower learning index for the optimal reach trajectory. In addition, we fully described the features underlying learning, including categorizing the first attempt during trials, failed reaches where mice make contact with the reward, the number of trials where no attempts were made, as well as how the pattern of these different behaviors varies in *Fmr1* KO mice. Our findings provide an essential framework for linking specific behavioral impairments in motor learning to the cellular and circuit mechanisms that support them.

## Introduction

One in 5000 males or one in 4000-8000 females have Fragile X Syndrome (FXS)^1–4^, yet therapies are limited, largely because our understanding of FXS is also limited. FXS is the largest single-gene cause of autism spectrum disorder (ASD) and inherited intellectual impairment, causing 1-2% of all ASDs^1,5–7^. FXS arises due to the loss of Fragile X Messenger Ribonucleoprotein (FMRP, encoded by the *FMR1* gene) usually because of a trinucleotide CGG repeat expansion in the promoter region of *FMR1*. FMRP is widely expressed in the brain and is involved in translational control of a large number of mRNAs, many of which are critical for synaptic function and plasticity. Approximately 50-60% males and 20% females with FXS show symptoms of ASD, and intellectual deficits are widespread^1^. The *Fmr1* knockout (KO) mouse is a well-established model of FXS, which replicates several of the signature features of human FXS: abnormal dendritic spine structure, altered synaptic plasticity, and behavioral deficits^1,8,9^. Alterations in synaptic plasticity have been demonstrated in several brain regions of the *Fmr1* KO mouse^10–13^.

Rodent models of FXS, including the *Fmr1* KO mouse, have permitted great strides in understanding the circuit, cellular, and molecular mechanisms underlying behavioral deficits in FXS. However, the ability to map neural functions to behavior requires a detailed knowledge of the different aspects of behavior in the rodent model. The behavioral features of FXS, and of ASDs more generally, are often difficult to replicate fully in rodents, given species-specific and ethologically relevant aspects of complex behaviors. In this context, motor function is a particularly appropriate and highly tractable behavior to investigate in rodents, allowing precise measurement and quantification of deficits.

Motor impairment is an important facet of the developmental problems in FXS and in ASD more generally^14–16^, although it has received less attention than other neurological deficits. However, motor skills are critical for normal development as well for interaction with other individuals and with the physical world. From a diagnostic standpoint, motor phenotypes are one of the first signs of impaired development in FXS^17^. Moreover, the severity of motor phenotypes diverges between children with both FXS and ASD and those with FXS without ASD^14^, fine motor skills are associated with social communication skills^15,18^, and motor deficits correlate with autism phenotypes and PDD-NOS (Pervasive Development Disorder – not otherwise specified)^19^. Therefore, given the prevalence and importance of deficits in motor behavior, understanding the neural basis of these deficits is essential for developing strategies to improve or alleviate them in FXS.

However, detailed descriptions of motor deficits in rodent models of Fragile X syndrome remain limited^20^. Although there is evidence that reflexive eyeblink conditioning is impaired^21^, our understanding of more complex forms of goal-directed motor learning remains incomplete. In turn, this lack of understanding limits our ability to link cellular/circuit deficits to specific aspects of motor function or motor learning, which is essential for a deeper understanding of the behavioral consequences of FXS. A key aim of therapeutic strategies in mouse models is to reverse the behavioral phenotype, highlighting how essential it is to fully describe the behavioral phenotype in the rodent model where cellular/molecular approaches for treatment are first described. In addition, animal and human behavior is complex and multifactorial, yet is often reduced to a single simple score. In contrast, the ability to link cellular and circuit function to behavior necessitates a precise and detailed analysis of behavior in mouse models.

In order to address this gap in knowledge, we investigated a form of goal-directed forelimb reach learning, in which several brain areas are involved, including the cerebellum^22^ and the motor cortex^23^. In particular, the cerebellum has been strongly implicated in Fragile X syndrome, both in humans and in the mouse model^21,24,25^, but deficits in the motor cortex are also likely to be involved in FXS-related deficits^26^. In addition to impaired gross and fine motor abilities, individuals with FXS demonstrate a hyperactivity phenotype, repetitive behaviors, and learning and memory impairments^1^, which can play a role in learning deficits in this task. Moreover, it has previously been demonstrated that *Fmr1* KO mice show deficits in this particular task^26^, making it an ideal paradigm for our investigation.

Here, we characterized in detail deficits in forelimb reach learning in the *Fmr1* KO mouse. First, we described that an overall deficit in learning was highly dependent on the experimental conditions, and could be alleviated by the presence of a cue. Next, by manually scoring different features of the learning, we described a range of deficits. By quantifying a granular subdivision of categories of reach success and reach failure, we described the behavior in detail, highlighting that the overall reduction in success could be broken down into specific deficits in learning. Finally, we used markerless tracking^27^ in order to map reach trajectories throughout learning and demonstrated that, while *Fmr1* KO mice are capable of significant improvement of reach trajectories over the time course of learning, they showed learning deficits.

## Results

Goal-directed forelimb reach learning was tested in *Fmr1* KO mice versus WT mice (see Methods for details). Briefly, mice were placed in a transparent box that had a slit on one wall, allowing access to a chocolate-flavored food reward that was placed on a platform outside the slit (Fig. 1 a-c). Mice had to learn to reach through the slit, grab the reward, and retrieve it back through the slit to their mouths. The learning process took 8 days, with mice given 50 consecutive learning trials per day. Both WT and *Fmr1* KO mice learned to perform this task, as measured by the percentage of successful trials within the 50-trial experiment on each day, over 8 days (Fig. 1 d, e). However, *Fmr1* KO mice performed significantly worse than WT mice, achieving a lower success rate (Fig. 1 d, e). This observation is consistent with previous experiments on a similar forelimb reach task^26^.

**Figure 1:**
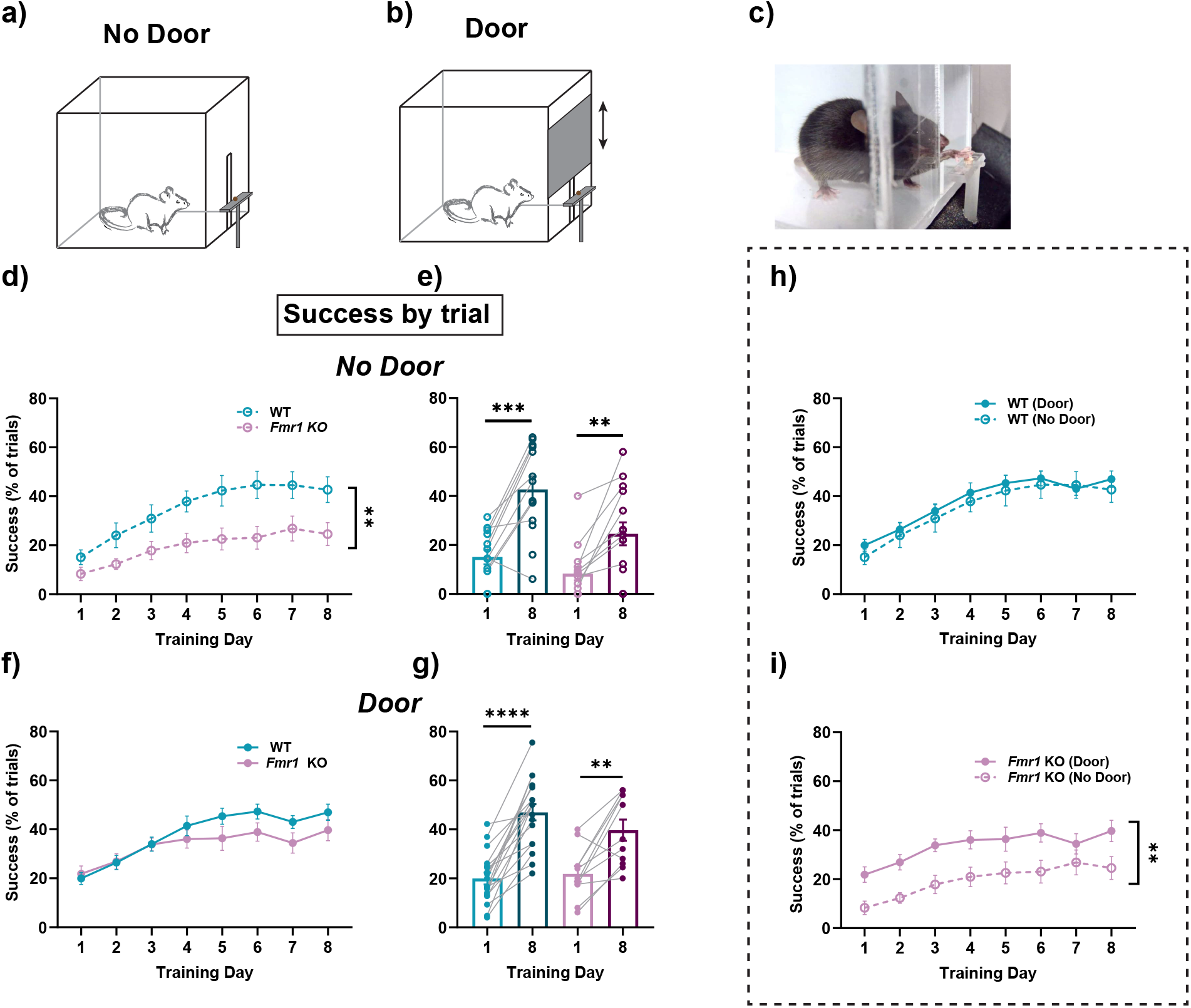
Forelimb reach learning was impaired in *Fmr1* KO mice. a) Schematic of experimental setup demonstrating behavior box and slit through which mouse had to reach to obtain a pellet reward; b) Schematic of experimental setup modified to add an automated door that opened at trial start and closed at trial end; c) Picture of mouse performing reach task; d) *Fmr1* KO mice showed less learning than WT mice did in the No Door condition: time course of forelimb reach learning. f) This deficit was alleviated in the Door condition. e, g) Comparison of Day 1 and Day 8 showed significant learning in both genotypes and in both conditions. Lighter colors indicate Day 1 and darker colors indicate Day 8. h, i) Same data as in d-g, comparing Door and No Door conditions and demonstrating that learning was significantly worse for *Fmr1* KO mice in the No Door condition. Time courses (d, f, h, i) were compared with a two-way ANOVA with repeated measures, or with a mixed-effects model. Paired comparisons (e, g) were done with a paired t-test or Wilcoxon matched pairs signed rank test. **** p<0.0001 ***p=0.001, **p<0.01.

We refined the task by the addition of an automated door that defined the trial structure (Fig. 1 b) (see Methods). Each trial now started with the door opening, allowing the mouse access to the pellet for a defined period of time before the door closed again. This refinement allowed the timing of each trial to be precisely controlled. Surprisingly, with the addition of this cue that signaled and defined the trial start, the performance of *Fmr1* KO mice was improved, and they no longer performed worse than WT mice (Fig. 1 f, g). Thus, the specific parameters of the task were critical for behavioral performance, and some feature of the door-cue alleviated part of the deficit in *Fmr1* KO mice (Fig. 1 h, i).

However, it has been well established that both individuals with Fragile X syndrome, and the *Fmr1* KO mouse, have deficits in motor function and in motor learning. Therefore, we investigated this apparent alleviation of the deficit further. We reasoned that defining success by trial number neglects information about the actual number of reach attempts made by a given mouse during learning, which is a critical factor in motor learning. Thus, instead of defining success as the percentage of successful trials, we counted the number of reach attempts during each trial, and measured the percentage of successful reaches out of the total targeted reach attempts (we defined targeted reaches as reaches that were made towards the food reward, in contrast to vain reaches, which were made when there was no reward present at all). All following results consider different parameters of learning in terms of the percentage of reaches. When analyzed in this manner, we observed that *Fmr1* KO mice do indeed have a deficit in learning (Fig. 2 a-d).

**Figure 2:**
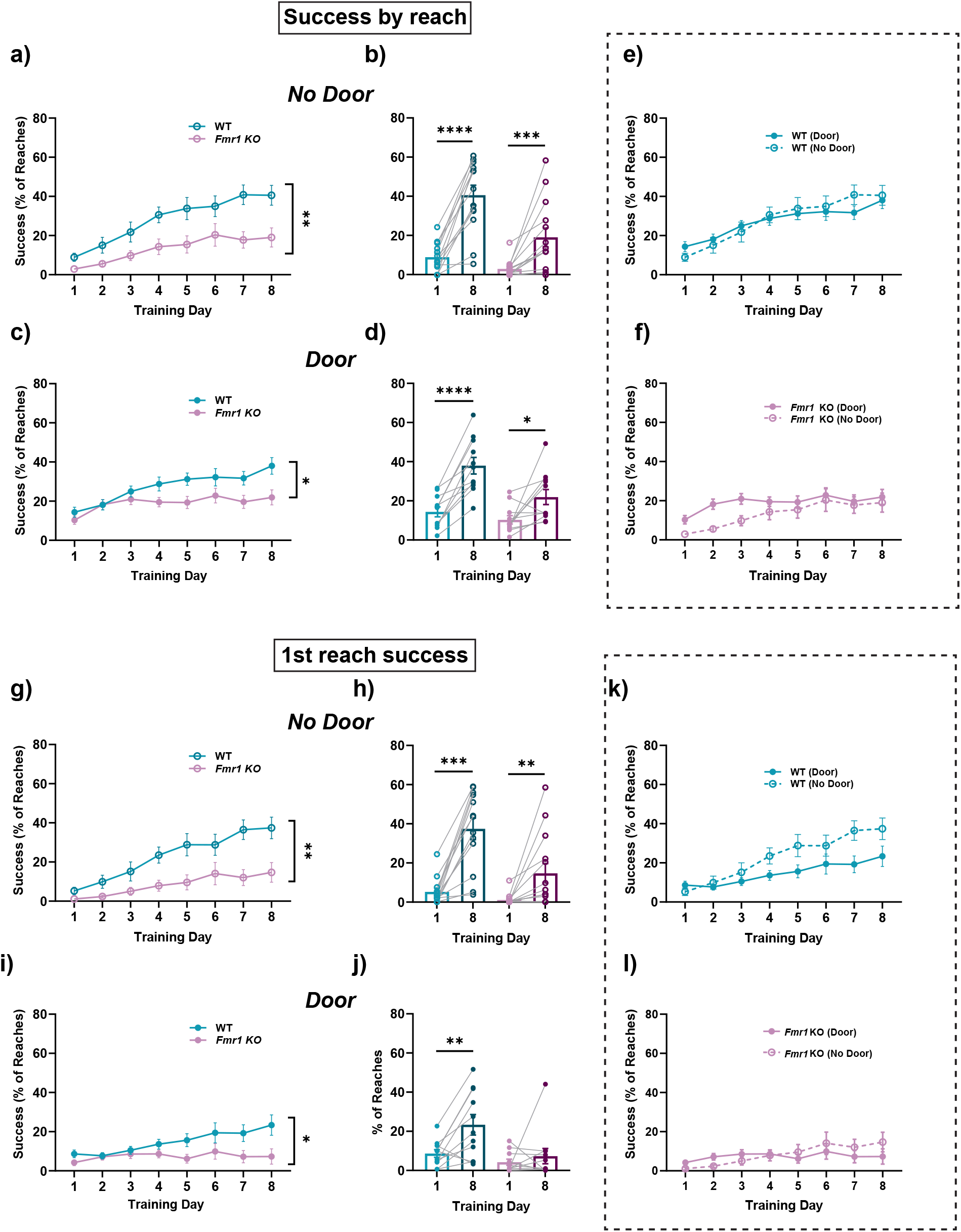
Forelimb reach learning was impaired in *Fmr1* KO mice when measured as a percentage of reaches. a, c) *Fmr1* KO mice showed significantly impaired learning in both the No Door and the Door condition. b, d) Both genotypes demonstrated significant learning from Day 1 to Day 8, under both conditions. Lighter colors indicate Day 1 and darker colors indicate Day 8. k, l) Same data as in a-d, comparing Door vs. No Door conditions. When success was measured as a percentage of reaches, there was no difference between the two conditions. g, i) The success of the first reach within a trial was also impaired in *Fmr1* KO mice. h, j) Both genotypes improved the rate of successful first reaches in the No Door condition, but only WT mice significantly improved in the Door condition. Lighter colors indicate Day 1 and darker colors indicate Day 8. k, l) Same data as in g-j. There was no statistical difference between the Door and No Door conditions for both genotypes. Time courses (a, c, e, f, g, i, k, l) were compared with a two-way ANOVA with repeated measures, or with a mixed-effects model. Paired comparisons (b, d, h, j) were done with a paired t-test or Wilcoxon matched pairs signed rank test. **** p<0.0001 ***p=0.001, **p<0.01* p < 0.05.

Success and failure during forelimb reach learning can come about because of many different reasons. Within this one metric of success or fail are hidden several possibilities: that the paw did not reach the target at all; that it did reach the target but that grasping and retrieval were impaired; that it took several consecutive reaches within a trial in order to refine the movement to obtain the target versus having a successful reach, grasp and retrieval on the first attempt within a trial; or even that the mouse was distracted and just made fewer attempts. We tried to disambiguate some of these possibilities by a detailed manual analysis of each reaching movement. Because the experiment and the analysis were performed by the same person (LFY), they were not blind to condition. Therefore, two additional scorers (RZ and AD), who were blind to the genotype of the mouse, repeated this analysis (Fig. S1 a, b), and obtained the same result.

First, we considered reach success both in the with-door and the without-door experimental condition. In both cases (Fig. 2 a-d) there was a significant impairment in learning in *Fmr1* KO mice, although they started on Day 1 without an impairment. The door and no-door conditions were not significantly different from each other (Fig. 2 e, f).

Next, we assessed whether a mouse had to make several reach attempts within a trial in order to successfully obtain the reward. During these multiple sequential attempts, mice can improve the trajectory of their reach and get closer to the target. We observed that in both the no-door condition (Fig. 2 g, h) and the door condition (Fig. 2 i, j), *Fmr1* KO mice showed a deficit in “1^st^ reach success”. This demonstrated that they were less able to retrieve the reward on the first attempt within a trial, highlighting an important aspect of the learning deficit. As previously, there was no significant difference between the no-door and door conditions (Fig. 2 k, l).

Another feature of learning that is encompassed within the success rate metric is the precise way in which the reach was not successful. In particular, it was recorded as a failure both when the mouse completely failed to make contact with the reward, defined as a “Complete Miss”, or when the mouse made a forelimb movement whose trajectory was appropriate enough to contact the reward but then failed to grasp or retrieve it, which we defined as a “Contact Miss”. It is important to distinguish whether the learning deficit in *Fmr1* KO mice arises because of an aberrant reach trajectory or because of an inability to learn how to grasp and retrieve it. Therefore, we measured complete misses under both the no-door (Fig. 3 a, b) and the door (Fig. 3 c, d) conditions, and demonstrated that there was a significant impairment in *Fmr1* KO mice. It is noteworthy that the deficit was more pronounced in the no-door condition. Indeed, neither WT nor *Fmr1* KO mice displayed significant learning in the door condition (Fig. 3 d). However, learning trajectories were not significantly different between the door and no-door conditions (Fig. 3 e, f). We speculate that the presence of the door makes the task easier by providing some guidance to the mouse’s paw. This is possible because mice attempt to reach the reward as soon as the door begins opening, which means that the bottom of the door is still sliding upwards when their paw reaches through the slit, and the paw often rests against the door edge. Therefore, they cannot make completely mistargeted reaches in the large open space above the reward, which we often observed in the early stages of learning when there was no door.

**Figure 3:**
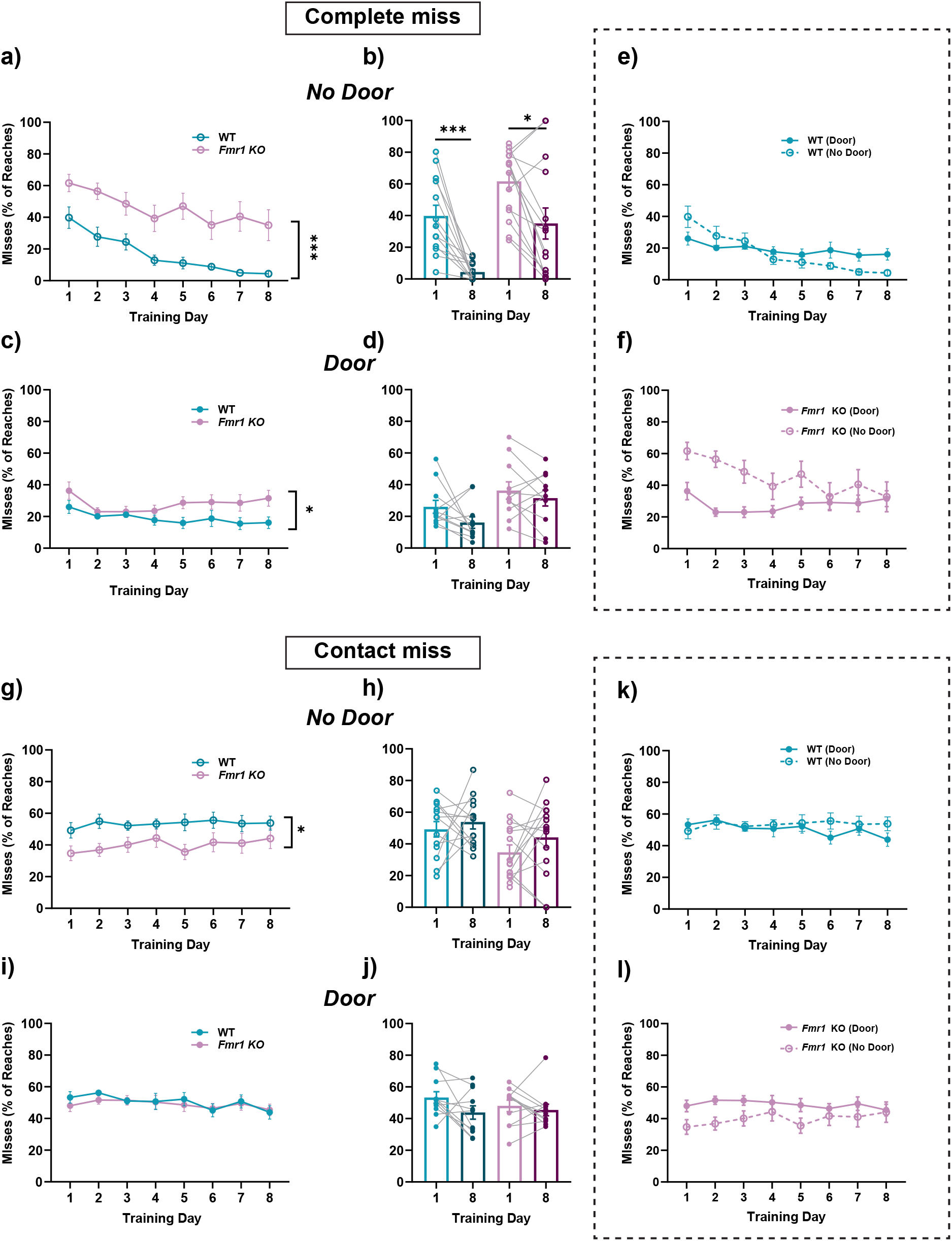
Learning was impaired in *Fmr1* KO mice when measured as a reduction in failures. a, c) In both No Door and Door conditions, *Fmr1* KO mice were impaired in terms of the percentage of complete misses they made. b, d) Both genotypes showed significant improvement of this metric only in the No Door condition, and not in the Door Condition. Lighter colors indicate Day 1 and darker colors indicate Day 8.e, f) Same data as in a-d, comparing Door vs. No Door, demonstrated no significant difference between the conditions. g, i) *Fmr1* KO mice showed a lower number of contact misses only in the No Door condition. h, j) In neither condition was there significant learning for either genotype. Lighter colors indicate Day 1 and darker colors indicate Day 8. k, l) Same data as g-j, comparing Door vs No Door conditions demonstrated no significant difference between the genotypes. Time courses (a, c, e, f, g, i, k, l) were compared with a two-way ANOVA with repeated measures, or with a mixed-effects model. Paired comparisons (b, d, h, j) were done with a paired t-test or Wilcoxon matched pairs signed rank test. **** p<0.0001 ***p=0.001, **p<0.01* p < 0.05.

In order to categorize the second type of miss category defined above, we tested if there were differences in learning in terms of reduction of contact misses, where the mouse’s reach successfully makes contact with the target but it either fails to grasp it or drops it before retrieval into the box. Again, there was a significant difference between WT and *Fmr1* KO mice, but this was only observed in the no-door condition (Fig. 3 g) and not in the door condition (Fig. 3 i). The observation that *Fmr1* KO mice had *fewer* contact misses than WT mice might be a reflection of their making more complete miss errors. In neither condition was there a significant improvement over the time course of learning. This suggests that this particular aspect of the reach was not what mice are improving on, but rather the ability to make an appropriate reach trajectory to the reward. In turn, this observation led to our experiments on analysis of the reach trajectory (Fig. 6). Although the door and no-door conditions showed markedly different results, a direct comparison between them over the course of learning (Fig. 3 k, l) did not attain significance.

Distinct from the actual reach trajectory, the grasp, and the retrieval are two further aspects of the learning task. While observing mouse behavior during this task, we noticed that mice sometimes continued to reach even when there was no reward present. Since the box was transparent, and the reward pellet was placed directly in front of the slit, it should have been visible to the mice. Yet, they sometimes kept trying to reach regardless. We called such reaches “Vain reaches”. Given that there is a tendency for repetitive behaviors in FXS^28^, we wondered whether *Fmr1* KO mice showed either more vain reaches, or less reduction in vain reaches over the time course of learning. Vain reaches were quantified as a percentage of total reaches (Fig. 4 a-f). Notably, there was a reduction of vain reaches, i.e. an improvement over the time course of learning, only in the condition with the door (Fig 4 c, d) and not in the no-door condition (Fig. 4 a, b). Direct comparison of the door and no-door conditions (Fig. 4 e, f) demonstrated a significant difference in WT mice, although *Fmr1* KO mice showed a trend that was not significant. Therefore, this was yet another aspect of the learning task that is markedly different between the two experimental conditions we tested, with the presence of the door cue allowing animals to make fewer vain reaches over the time course of learning.

**Figure 4:**
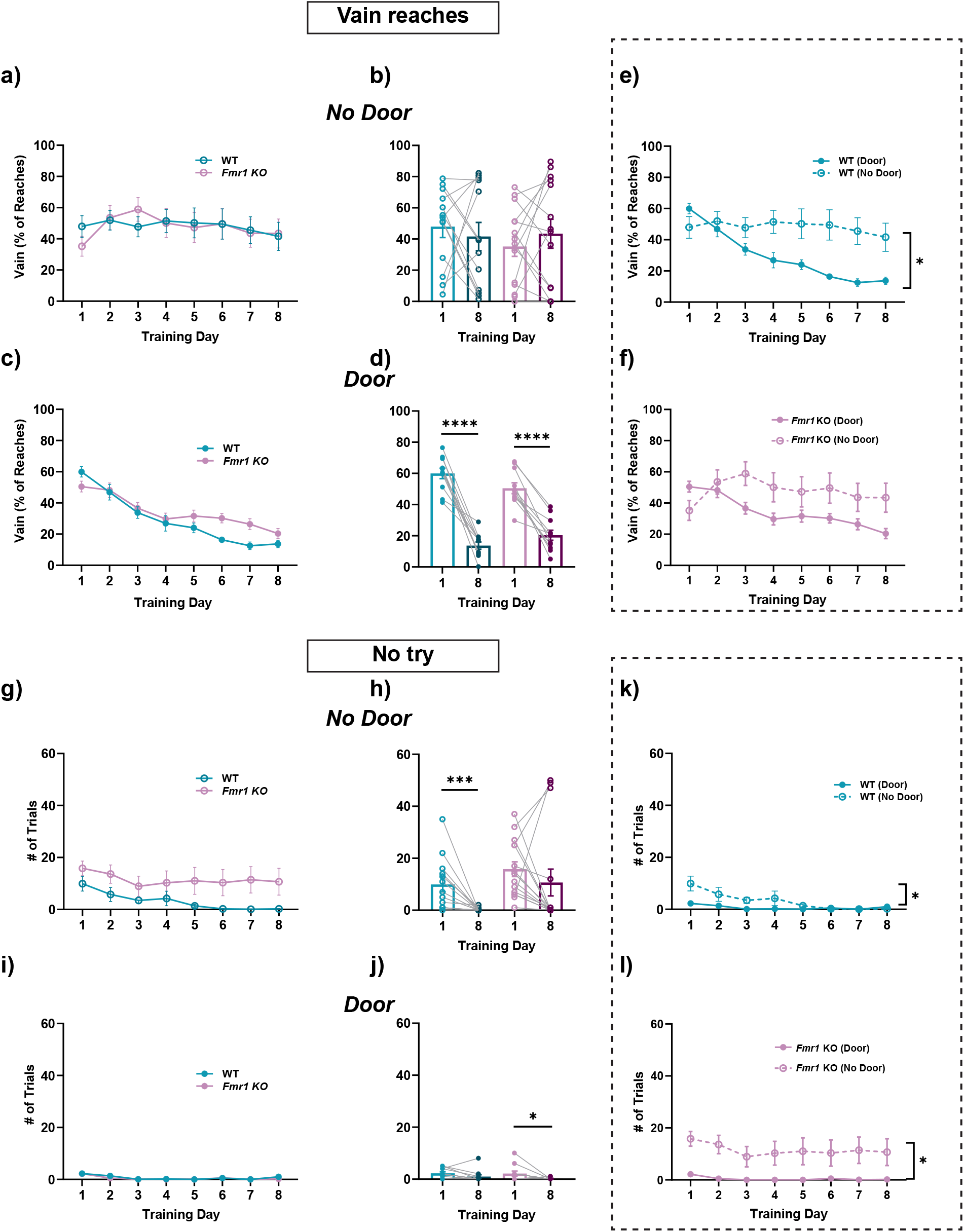
Vain reaches and no tries did not explain lower success rates in *Fmr1* KO mice. a, c), In both Door and No Door conditions, there was no difference in the time course of vain reaches as a percentage of total reaches. b, d) However, both genotypes learned only in the Door condition, and not in the No Door condition. Lighter colors indicate Day 1 and darker colors indicate Day 8.e, f) Same data as in a-d, replotted to compare Door vs. No Door conditions. WT mice showed significantly different improvement in the percentage of vain reaches in the Door condition, compared to No Door. g, i) There was no significant difference in the time course of trials where the mouse made no tries, in both No Door and Door conditions. h, j) There was significant improvement in WT in the No Door condition and in *Fmr1* KO in the Door condition. Lighter colors indicate Day 1 and darker colors indicate Day 8. k, l) Same data as in g-j, plotted to compare Door vs No Door conditions across genotypes. Both genotypes showed significantly different behavior with the two conditions. Time courses (a, c, e, f, g, i, k, l) were compared with a two-way ANOVA with repeated measures, or with a mixed-effects model. Paired Young et al. 2024 (preprint) 9 comparisons (b, d, h, j) were done with a paired t-test or Wilcoxon matched pairs signed rank test. **** p<0.0001 ***p=0.001, **p<0.01* p < 0.05.

In addition, we noticed that *Fmr1* KO mice appeared to miss several trials just because they were distracted and were in another part of the box, or only noticed that the reward pellet for the next trial was in position shortly before the end of the trial. We therefore defined another metric called “No try”, and counted the number of trials where the animals did not make reach attempts. As expected, this number was much higher in the no-door condition (Fig. 4 g, h) than in the door condition (Fig. 4 i, j). It is possible that this was due to the inherent sound or visual cue of the door opening, which allowed animals to know when to pay attention because a new trial was starting. Although *Fmr1* KO mice did not show a significant difference in the time course of learning when compared to WT (Fig. 4 g), WT mice showed a significant improvement, while *Fmr1* KO mice did not (Fig. 4 h). In addition, this effect appeared to be lowered and potentially reversed in the door condition. First, there were very few no tries for both genotypes, but there was a significant improvement in *Fmr1* KO mice and not in WT. However, the total number of no tries was so small that the effect of noise is considerable. A direct comparison of door and no-door conditions showed a significant difference for both genotypes, highlighting yet another important effect of the door cue on learning behavior.

In order to provide an overview of these results, we plotted the different behaviors at the end of learning as pie charts. The categories of success, complete miss, and contact miss form the majority of reaches. The differences between the varied aspects of learning described above are apparent (Fig. 5 a-d), where WT mice showed many more successes than *Fmr1* KO under both conditions. However, the distribution of types of failure varied depending on condition.

**Figure 5:**
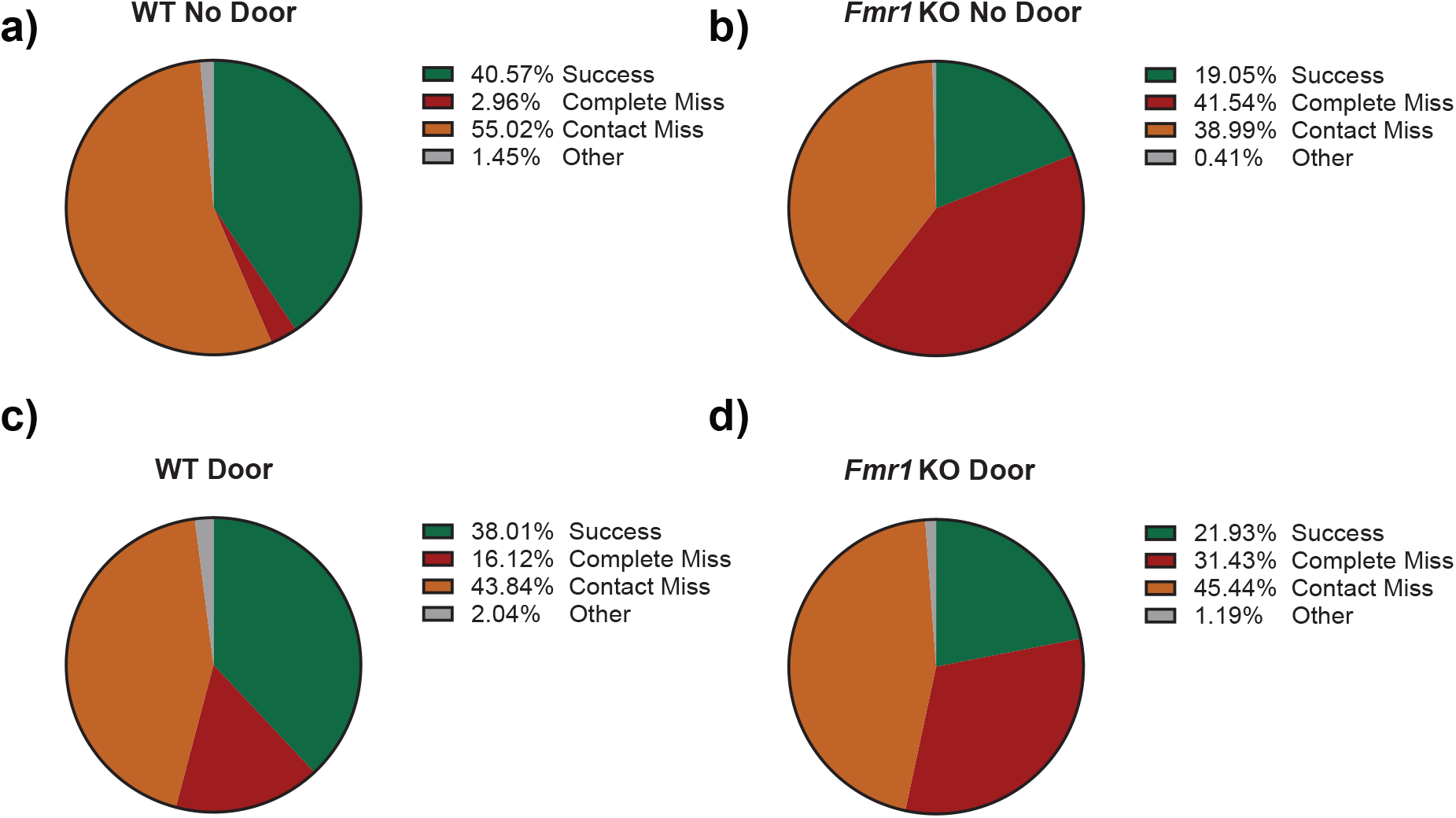
Distribution of behavioral outcomes varies between *Fmr1* KO and WT mice on the final day of learning. a-d) Successes, complete-misses, contact-misses, and “other” categories of reach, across both genotypes and door conditions.

In patients with Fragile X syndrome, due to the X-linked nature of the deficit, females often show milder phenotypes^1^. However, in the mouse model, FMRP is absent in both sexes and the degree of X-inactivation does not play a role. Notwithstanding this difference between the mouse model and humans, sex differences have been described in the *Fmr1* KO mouse^29^, but also see^30^. Therefore, in order to determine whether the sexes show differences in forelimb reach learning, we separated males and females and compared their behavior (Fig. S2 a-d). We found no significant difference between the sexes, in either genotype, and in both door and no-door conditions.

Since our results above suggested that there was a difference in the reach trajectory between *Fmr1* KO and WT mice, we went on to measure these trajectories using a well-established, markerless, deep-learning based method for pose estimation, DeepLabCut^27^. Over the 8 days of learning, the reach trajectory was strikingly refined, in both WT and *Fmr1* KO mice (Fig. 6 a). We quantified this refinement in several ways. First, we measured the distance for each reach in pixels (Fig. 6 b). In order to better understand the process of refinement, we also defined a Learning Index as the difference in any metric between Day 1 and Day 8. Although both WT and *Fmr1* KO mice showed a reduction in the distance per reach, there was no difference between them (Fig. 6 b, c). Next, we compared the x and y dimensions of the reach, in pixels. We noticed that *Fmr1* KO mice made reaches that were more spread out in the y-axis, i.e., they were further away from the direct path to the target. We quantified this as ΔY, which was defined as the difference between the maximum y coordinate and the minimum y coordinate for each reach. As expected, ΔY was also reduced over learning (Fig. 6 d, e). Although there was no significant difference in the time course plots in Fig. 6 d, the learning index was significantly different between WT and *Fmr1* KO (Fig. 6 e). This suggests that KO mice were less able to refine their trajectories to reach the target efficiently. We also measured a metric we defined as Max X. Since the extent of the reach is defined at one end by the slit, variability in the reach comes from how far from the slit the reach trajectory extended, often overshooting the target. As shown in Fig. 6 a, as learning progressed, the reaches got shorter and no longer overshot the target. This was apparent in the reduction in Max X (Fig. 6 f, g). However, there was no difference between the genotypes. Overall, our analysis of the reach trajectory showed that although *Fmr1* KO mice did successfully refine their reach over learning, there were nevertheless impairments in the extent of the reach and the ability to refine the reach over learning.

**Figure 6:**
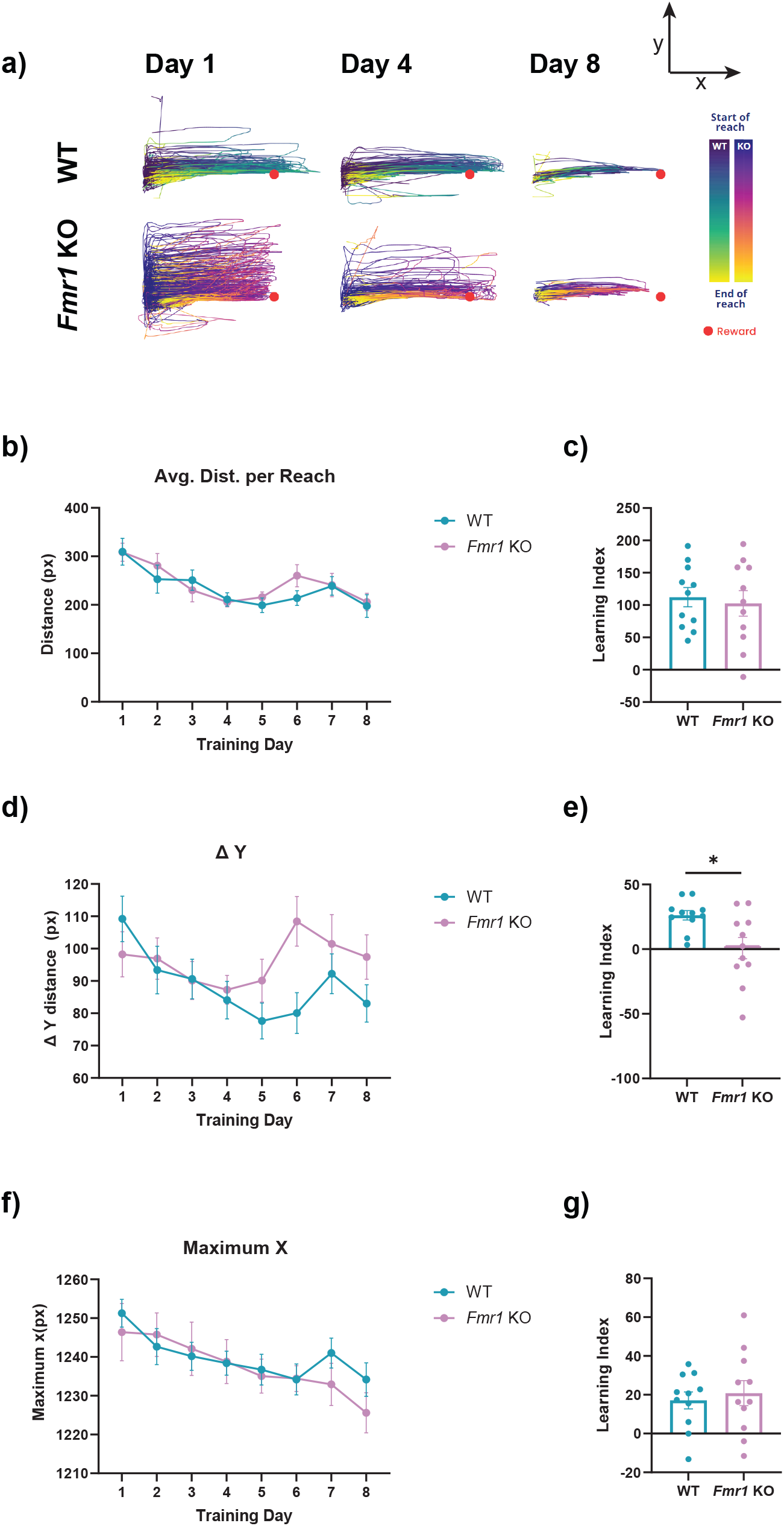
*Fmr1* KO mice demonstrated improvement in reach trajectory over learning, but deficits in learning. a) Example traces of reach trajectories over learning for individual example mice of each genotype. The reach is tracked from the slit at the left towards the target, marked in red, on the right b) The time course of average distance per reach was not different between *Fmr1* KO mice and WT mice and c) neither was the Learning Index related to this metric. d, e) There was a reduction in the ΔY distance over learning, measured as the difference between the maximum y position and the minimum y position for each reach, only in WT mice. This was reflected in a significantly lower learning index in *Fmr1* KO mice. f, g) improvement in the Max X, which is the maximum extent of the reach in the x dimension, was not different between *Fmr1* KO and WT mice.

## Discussion

Human patients with FXS demonstrate a range of motor impairments^20,28,31^. Motor learning is ideal for investigation in animal models, in comparison to more complex human behaviors that are difficult to replicate in mice. In addition, motor learning can be precisely measured and quantified, raising the possibility of linking specific motor deficits to their underlying cellular or circuit basis. Therefore, in order to determine motor learning deficits in the *Fmr1* KO mouse, we compared a goal-directed reach learning task between *Fmr1* KO and WT mice.

We demonstrated that *Fmr1* KO mice were capable of a form of goal directed forelimb reach learning in which mice had to learn to retrieve a reward pellet through a narrow slit. In alignment with previous results, we demonstrated an impairment in this forelimb reach learning in *Fmr1* KO mice, when compared to WT mice^26^. This impairment was quantified as an overall success rate. Intriguingly, we discovered that the addition of an automated door that signaled the start and end of the trial, and thereby provided a cue as to when the reward was available, alleviated some specific aspects of the deficit in *Fmr1* KO mice. In particular, when successes were quantified as a percentage of the total trials, there was no longer a significant difference between *Fmr1* KO and WT mice. However, more detailed analysis did demonstrate that there was still an impairment in motor learning. It is possible that the behavioral changes due to a door may have to do with attention deficits, with the door providing a cue signaling availability of the reward. When the door cue was absent, we observed that *Fmr1* KO mice were often in a different part of the box and were not paying attention when a trial started with the placement of the food reward. Thus, the door may provide a signal to pay attention. This may be particularly relevant to FXS because in human males with FXS, attention was found to be impaired^32–34^. There is also evidence that males with FXS can maintain attention for short periods of time (a few min.), but not for longer periods of sustained attention^35^. It is also possible, as described in the Results section, that the physical presence of the door constrains early reaches while the door is opening, and thereby prevents mistargeted reaches in the space above the reward pellet. Thus, our data also showed that the parameters of learning were critical to the ability to fully describe motor deficits in *Fmr1* KO mice, and that the exact experimental conditions made a large difference to the outcome.

Learning in goal-directed reach tasks is often described by a single success score. However, this single score encompasses a range of different parameters of learning that could be responsible for an overall lower success rate. Therefore, in order to identify what these specific parameters or features might be, we performed a more granular analysis of the reasons behind the reduced success rate in *Fmr1* KO mice. First, since a mouse can make several reaches within a trial, we analyzed how success varied when scored as a percentage of trials, or as a percentage of targeted reaches. Our results demonstrated that the outcomes of these two different analyses were different, which was also apparent when comparing the door and no door conditions. Next, out of the multiple reach attempts a mouse could make within a trial, our analysis demonstrated that *Fmr1* KO mice were less successful at the first reach within a trial. Furthermore, the pattern of failures they made, i.e., entirely missing the target, or making contact but failing, provide insight into different aspects of the motor deficit. Future experiments will be required to further clarify the distinctions between deficits in reach trajectories, reach kinematics, or grasping and retrieval. We also determined that, at least under the specific conditions of this task, *Fmr1* KO mice did not make a higher percentage of “vain reaches” than WT, which might have been expected given the prevalence of repetitive behaviors in ASD. In addition, depending on whether there was a door cue or not, *Fmr1* KO mice also had more trials in which they did not make attempts.

In addition to manual analysis, we performed markerless tracking analysis using DeepLabCut. Over the time course of learning, the reach trajectory was refined in both WT and *Fmr1* KO mice, reflected by a reduction in the average distance of each reach. However, *Fmr1* KO mice made reaches that were not as optimally directed to the target, reflected by a greater vertical extent of each reach and measured as a ΔY. These results suggest that *Fmr1* KO mice were not able learn optimally targeted reaches. Our current study has the caveat that our field of view was limited by the camera angle and resolution, and did not permit accurate analysis of the grasping movement of the digits. Moreover, we used the 2D version of DeepLabCut; extending our analysis to 3D DeepLabCut^27,36^ is an important future goal. Therefore, a more detailed analysis may highlight aspects of the reach that we were not able to measure. Nevertheless, even with these limitations, we were able to determine how the trajectory improved over time, and how this improvement was impaired in *Fmr1* KO mice.

The relevant brain regions, underlying molecular mechanisms, and potential therapeutic targets in Fragile X syndrome have long been a focus of intense interest^6,10,37–4243^. Various forms of behavior and learning have also been previously studied in rodent models of FXS^9,26,30,44–46^. However, the ability to capitalize on studying mouse models for preclinical studies is contingent on a deeper understanding of the behavioral deficits they demonstrate, and their neural basis. There is a diversity of behavioral impairments in Fragile X syndrome, ranging from attention deficit hyperactivity disorder, to intellectual disability, to difficulties with social behavior. However, complex cognitive tasks are difficult to model accurately in mice without some degree of anthropomorphism. In addition, linking complex behaviors to their underlying cellular or circuit-level substrates remains challenging. In contrast, motor learning is an ideal system both in terms of its relevance to mice, and the ability to precisely measure and quantify it. Such precise measurement and quantification provides the basis for linking behavioral features to cellular targets, particularly in the cerebellum and the motor cortex^22,26^. Thus, our investigation into motor learning deficits in *Fmr1* KO mice provides the framework for more detailed investigation of the underlying intracellular, cellular, and circuit mechanisms of Fragile X syndrome. Overall, our description of deficits in forelimb reach learning in *Fmr1* KO mice brings us closer to being able to link specific features of behavioral dysfunction to their neural correlates.

## Acknowledgements

We thank Dr. Arnold Hayer for his comments and suggestions. A.S. received funding from the Canadian Institutes of Health Research (CIHR) Project Grant PJT-178281, Natural Sciences and Engineering Research Council of Canada (NSERC) Discovery Grant RGPIN-2020-07073, Canada Foundation for Innovation John R. Evans Leaders Fund (CFI-JELF) Equipment Grant 38053, The Scottish Rite Charitable Foundations, Fonds de recherche du Quebéc – Santé (FRQS) Chercheurs Boursiers/ Chercheuses Boursières, FRQS Établissement de jeunes chercheurs, a New Recruit Start-Up Supplement from Healthy Brains for Healthy Lives (HBHL) and the Canada First Research Excellence Fund (CFREF), and startup funding from the Research Institute of the McGill University Health Centre.

## Author contributions

Experiments were designed by AS and LY, experiments were conducted by LY, analysis was performed by LY, AD, RZ, and AS. The manuscript was written by AS and LY.

## Competing interest statement

The authors declare no competing interests.

## Materials and Methods

### Mice

All experiments were performed in accordance with the policies of the Canadian Council on Animal Care, using protocols approved by the Montreal General Hospital Facility Animal Care Committee, using mice of both sexes aged P55-P93. Mice were either wild type (WT), C57BL/6J (Strain # 000664, The Jackson Laboratory), or *Fmr1* KO (Strain # 003025, The Jackson Laboratory). Genotyping was performed following the protocol provided by the Jackson Laboratory. Mice were maintained on a 12-hour inverted day-night cycle with *ad libitum* access to food and water except when food restricted for learning, as described below.

### Forelimb reaching task

All mice were handled for 5 min. each day for 5 days before habituation to the behavior apparatus. Mice were food restricted to ∼85% of their original body weight starting 3 days prior to the first day of the experiment. The learning paradigm had three stages, “habituation” to introduce the mouse to the chocolate pellet food reward (Bio-Serv Dustless Precision Pellets, chocolate flavor) and learning environment, “shaping” to teach the mouse to use its paw to retrieve the food reward, and “training”, which was the actual learning phase of the task. On the first day during habituation, each mouse was placed into a custom-built clear plexiglass box (“behavior box”) with a vertical 5 by 9 cm opening (“slit”) in the front wall (Fig. 1 a-c) for 20 min. Food pellets were scattered in the box. Starting on the second day, mice had between 1-4 days of “shaping” to meet the requirements needed to pass onto the training stage. Each day of shaping was 20 min. in duration. Shaping involved placing a food pellet on a platform outside but close to the slit, within reach of the mouse’s tongue. Once the pellet was consumed, a new one was replenished and the distance of the pellet to the slit was incrementally increased (to a maximum of 9 mm). This taught the mouse that the pellet was rewarding, and introduced them to the idea of not using just the tongue to retrieve it. Then, one pellet was placed 1 cm away from the slit and the mouse had to make a single reach attempt towards it. The criteria to pass shaping were a minimum of 2 pellets eaten from outside the slit, a maximum of 3 successful pellet retrievals at any distance, and a maximum of 10 reach attempts overall. All mice had to pass shaping before they could proceed to the training phase.

On the first day of training, all mice received a “reminder trial” where the pellet was placed into a 4 mm wide indentation (“divot”) on a platform 1 cm away from the slit outside the behavior box. This trial was only given on the first day of training and did not count towards the trials used to analyze learning. Once the mouse made reach attempts that made contact with the pellet, the pellet was removed and the training day began. Mice underwent 8 consecutive days of training, and each training day consisted of 50 trials, where each trial was defined by a timed window where the mouse had access to the pellet before the pellet was removed. Mice had 13s of access to the pellet and 10s of wait time between trials in the No Door condition.

### Forelimb reaching task with automated door

On the first day of training, all mice received a “reminder trial” where the pellet was placed into a 4 mm wide indentation (“divot”) on a platform 1 cm away from the slit outside the behavior box. This trial was only given on the first day of training and did not count towards the trials used to analyze learning. Once the mouse made reach attempts that made contact with the pellet, the pellet was removed and the training day began. Mice underwent 8 consecutive days of training, and each training day consisted of 50 trials, where each trial was defined by a timed window where the mouse had access to the pellet before the pellet was removed. Mice had 13s of access to the pellet and 10s of wait time between trials in the No Door condition.

For additional cohorts of mice, including all with DLC-based reach tracking, an automated door was attached to the behavior box, which opened and closed at the same time intervals listed above to give mice the same amount of access time to the pellet as they did in the absence of the door. The protocol for the task remained the same, with the exception of including the activation of the automated door (14s open, 8s closed) during shaping days in order to familiarize mice to the sound and sight of the door.

### Manual reach scoring analysis

All behavioral experiments were recorded using a webcam at 60 frames per second (fps). Each reach was manually classified (Table 1). A reach was defined as any reaching movement from when the paw dorsum passed outward through the slit and returned through the slit. A successful reach was defined as one continuous reaching and grasping motion where the pellet was successfully retrieved and brought to the mouse’s mouth without letting the pellet touch the floor of the behavior box. A targeted reach was defined as any reach where the pellet is on the platform. If any mouse did not perform any valid reaches during a trial, the trial is marked as “No Try”. A trial was considered successful if there was one successful reach attempt during the trial. Success rate calculated by successful trials did not include trials with no reach attempts. or additional cohorts of mice, including all with DLC-based reach tracking, an automated door was attached to the behavior box, which opened and closed at the same time intervals listed above to give mice the same amount of access time to the pellet as they

**Table 1.**

| Reach Type | Description |
| --- | --- |
| 1<sup>st</sup> Reach Success | A successful reach made on the first attempt of the trial. |
| Complete Miss | A reach where the paw did not come into contact with the pellet. |
| Contact Miss | A reach where the paw came into contact with the pellet, but did not retrieve the pellet back past the slit. The pellet was not consumed during this trial. |
| Vain | Pellet was not on the platform when the reach was made, and was therefore not classified as a targeted reach. |

### 2D reach tracking analysis with DeepLabCut

An opaque plexiglass door that was added to the behavioral box was connected to a motor that was controlled by a Raspberry Pi running a custom Python script. The opening and closing of the door were synchronized to the activation and deactivation of the camera (OptiTrack Prime Color) used to record the mice. All recordings with this camera were made at an angled left side view at 240 fps, and every trial was recorded for each mouse over 8 days of learning the task. A subset of these videos was used to train a deep learning model to track the paws using DeepLabCut^27^.

#### DeepLabCut

All two-dimensional analysis was performed using DeepLabCut to track the paws and pellet. Over 2,500 frames were manually extracted from videos selected to maximize the diversity of mouse behaviors, and the left paw, right paw, nose, and pellet were manually labeled for each frame to train the network (ResNet-50). The “p-cutoff”, the likelihood threshold to help distinguish labels with a higher probability of accuracy from uncertain ones, was set to 0.85, and network performance had a 3.44 pixel (px) test error. Novel behavior videos were then inputted into DeepLabCut to extract pixel coordinate points of all labeled elements for further analysis.

#### Reach detection and extraction

Reach analysis and data filtering were conducted in Python. Data was filtered by set tracking bounds and a defined p-cutoff value. Individual reaches were then isolated from trial data using an algorithm. A reach was detected if it met a list of criteria: A reach must have at least 6 data points. If a datapoint and its subsequent 5 data points were within the tracking bounds, the vectors between each adjacent datapoint for 5 points succeeding the first were calculated 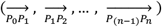. If at least 4 out of 5 vectors were towards the direction of the pellet, the first datapoint of the sequence was marked as the start of a reaching movement. This process was then repeated for the opposite direction to determine the point at which the paw began retracting towards the slit, marked as the reach endpoint. This was continuously repeated, reversing the direction of vector comparisons until all tracked reaches in the trial had been exhausted.

## Notes

### Competing Interest Statement

The authors have declared no competing interest.

