## Supplementary Figures for "Deficits in forelimb reach learning in a mouse model of Fragile X syndrome"

#### Supplementary Information

#### Supplementary Figures and Legends:

**Supplementary Figure S1:** Reach analysis was performed blind to genotype, and replicated the data in Figure 2. a, b) *Fmr1* KO mice showed significantly impaired learning in both the No Door and the Door condition.

**Supplementary Figure S2:** a-d) There was no significant difference between male and female mice in the time course of learning for both genotypes and for both Door and No Door conditions.

### Figure S1

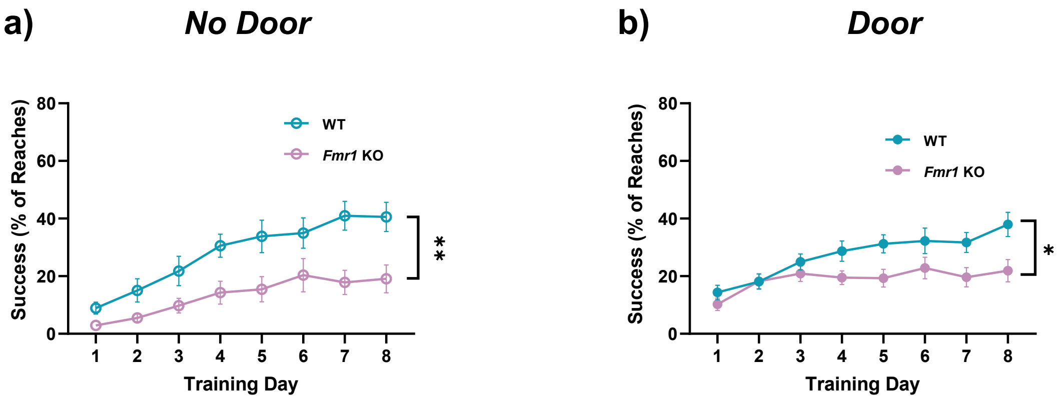
a)
No Door
Success (% of Reaches)
80
60
40
20
0
1 2 3 4 5 6 7 8
Training Day
WT
Fmr1 KO
**
b)
Door
Success (% of Reaches)
80
60
40
20
0
1 2 3 4 5 6 7 8
Training Day
WT
Fmr1 KO
*

#### Figure S2

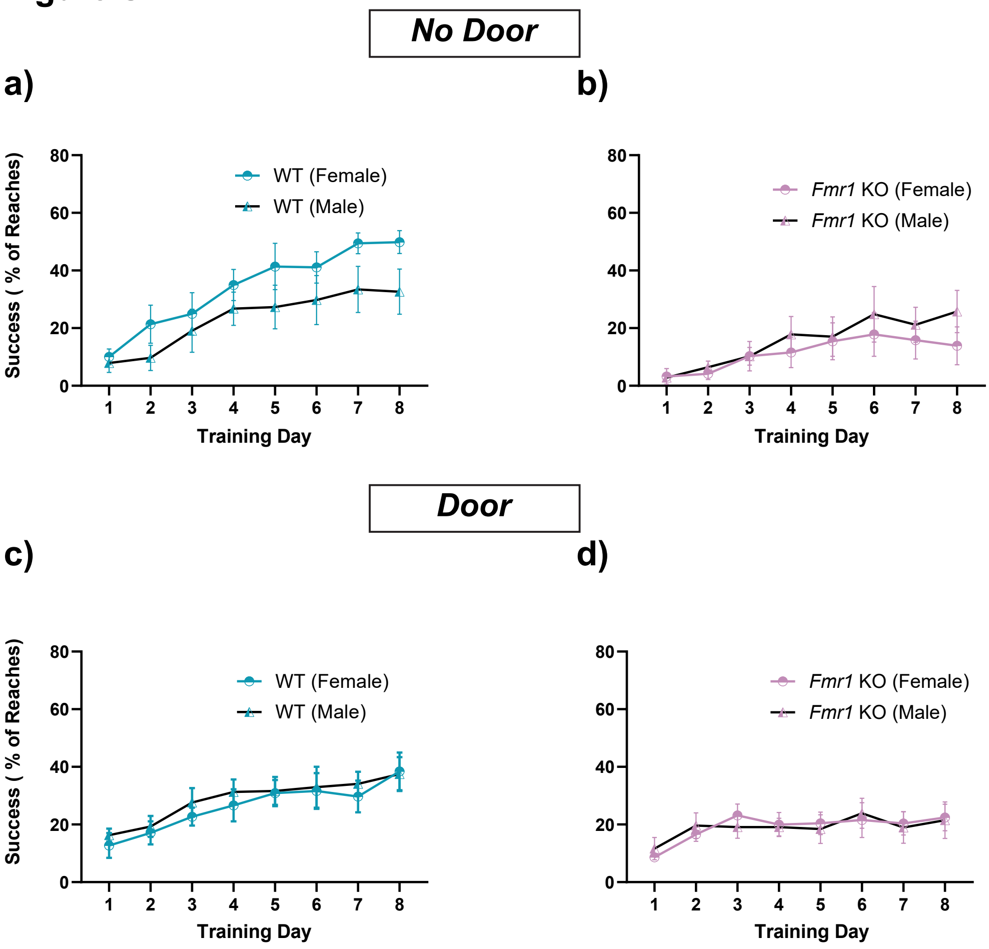

No Door
a)
b)
Door
c)
d)
Success ( % of Reaches)
Training Day
WT (Female)
WT (Male)
Fmr1 KO (Female)
Fmr1 KO (Male)
0
20
40
60
80
1
2
3
4
5
6
7
8
